# Shared features of blastula and neural crest stem cells evolved at the base of vertebrates

**DOI:** 10.1101/2023.12.21.572714

**Authors:** Joshua R. York, Anjali Rao, Paul B. Huber, Elizabeth N. Schock, Andrew Montequin, Sara Rigney, Carole LaBonne

**Affiliations:** Dept. of Molecular Biosciences, Northwestern University, Evanston, Il 60208; NSF-Simons Center for Quantitative Biology, Northwestern University, Evanston, Il 60208; New York University Langone Medical Center

## Abstract

The neural crest is vertebrate-specific stem cell population that helped drive the origin and evolution of the vertebrate clade. A distinguishing feature of these stem cells is their multi-germ layer potential, which has drawn developmental and evolutionary parallels to another stem cell population—pluripotent embryonic stem cells (animal pole cells or ES cells) of the vertebrate blastula. Here, we investigate the evolutionary origins of neural crest potential by comparing neural crest and pluripotency gene regulatory networks (GRNs) in both jawed (*Xenopus*) and jawless (lamprey) vertebrates. Through comparative gene expression analysis and transcriptomics, we reveal an ancient evolutionary origin of shared regulatory factors between neural crest and pluripotency GRNs that dates back to the last common ancestor of extant vertebrates. Focusing on the key pluripotency factor *pou5* (formerly oct4), we show that the lamprey genome encodes a *pou5* ortholog that is expressed in animal pole cells, as in jawed vertebrates, but is absent from the neural crest. However, gain-of-function experiments show that both lamprey and *Xenopus pou5* enhance neural crest formation, suggesting that *pou5* was lost from the neural crest of jawless vertebrates. Finally, we show that *pou5* is required for neural crest specification in jawed vertebrates and that it acquired novel neural crest-enhancing activity after evolving from an ancestral *pou3*-like clade that lacks this functionality. We propose that a pluripotency-neural crest GRN was assembled in stem vertebrates and that the multi-germ layer potential of the neural crest evolved by deploying this regulatory program.

## Main

Embryogenesis is a progressive restriction of developmental potential, beginning with a totipotent egg. In vertebrates, a key population of cells defies this this principle—the neural crest. Neural crest cells arise in the ectoderm at the neural plate border, yet they display a multi-germ layer potential greater than that of their ostensibly ectodermal progenitors^1–3^. These stem cells are central to the vertebrate body plan because they contribute diverse cell types, tissues, and structures that were essential to the origin and diversification of vertebrates, including cartilage, bone, and smooth muscle of the craniofacial skeleton, sensory neurons and glia of the peripheral nervous system, colorful patterns of pigmentation in skin, feathers, and scales, and primordia of the teeth and heart^3–5^.

Insights into the origins of neural crest potential came from the realization that these stem cells share GRN components with pluripotent stem cells of the vertebrate blastula^1,3,6,7^. Indeed, in *Xenopus* neural crest regulatory genes such as *snai1*, *zic1*, *id3*, *tfap2a,* and *foxd3* are co-expressed with the core pluripotency genes *sox2, pou5*, *myc*, and *ventx/nanog*^8,9^ in blastula (“animal pole”) stem cells, and are required for maintenance of pluripotency^1^. These and other shared features^10,11^ led us to hypothesize that the neural crest evolved by retaining molecular characteristics of those earlier cells. Under this model, selective retention of blastula-stage pluripotency enables neural crest cells to escape early lineage restriction and contribute a diverse array of cell types to the vertebrate body plan. In mouse embryos it was also found that core pluripotency factors are required for neural crest cell potential^12^. However, there it was suggested that pluripotency factors such as pou5 become re-activated—rather than retained—in neural crest progenitors, thereby reprogramming these cells to have expanded cellular potential^12^. By contrast, recent work in chick embryos reported that core pluripotency factors *pou5*, *nanog* and *klf4* share broad ectodermal expression from the epiblast through gastrulation and neurulation before becoming restricted to the neural crest, similar to that of *Xenopus*^13^.

These studies all point to the developmental potential of the neural crest in jawed vertebrates being linked to the deployment of pluripotency GRN components, and that a dual pluripotency-neural crest GRN may be evolutionarily conserved across vertebrates. However, demonstrating this requires investigating the relationship between pluripotency and neural crest GRNs in the other major clade of vertebrates—the jawless cyclostomes (lampreys, hagfish). To this end, we compared the pluripotency and neural crest GRNs in a jawless vertebrate, the sea lamprey, *Petromyzon marinus*, to that of a jawed vertebrate, the frog, *Xenopus laevis* (**Fig. 1a, b**).

**Fig. 1.**
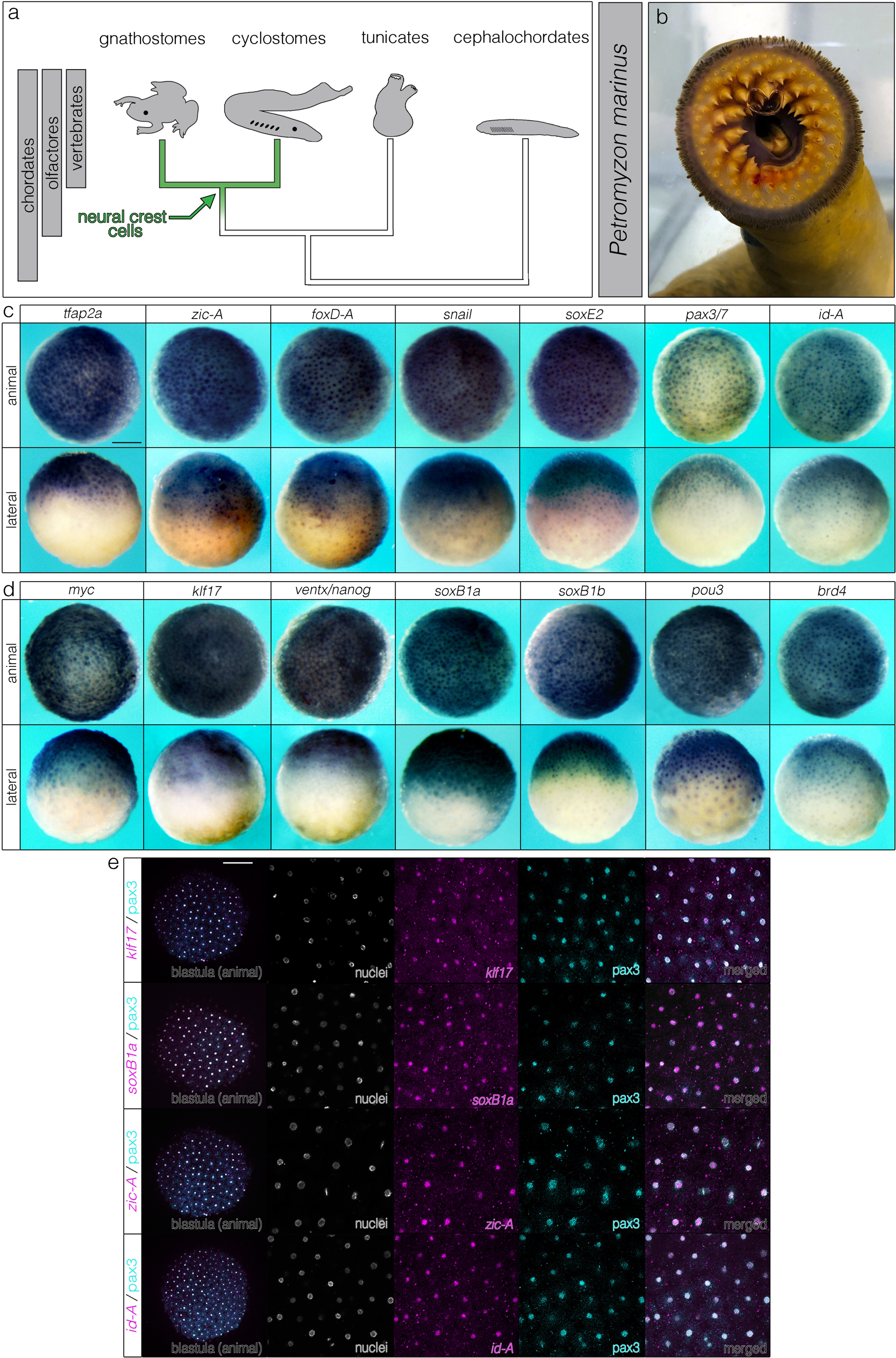
Lamprey animal pole cells co-express neural crest and pluripotency GRN components. (a, b) Phylogenetic framework for investigating the evolutionary origins of neural crest potential using the sea lamprey (*Petromyzon marinus*) and *Xenopus laevis*, species separated by 500 million years of evolution. (c) *in situ* hybridizations of canonical neural crest and (d) pluripotency genes in animal pole cells of blastula-stage lamprey embryos. (e) Neural crest and pluripotency factors co-localize in lamprey animal pole cells. Reproducible on n ≥ 10 embryos per time point for n ≥ 3 experiments. Scale bars: 250 μm.

## Results

### Neural crest and pluripotency GRN factors are co-expressed in the lamprey blastula

If neural crest and pluripotency GRNs share a common developmental and evolutionary origin, then a shared expression signature between these cells should be found in both jawed and jawless vertebrates. In *Xenopus*, this signature is revealed by co-expression of neural crest (*sox5*, *snai1*, *id3*, *tfap2a*, *foxd3*, *zic1*, *ets1*) and pluripotency (*sox2*, *sox3*, *ventx2.2*, *myc*, *pou5f3.2, pou5f3.3*) genes^1^ in blastula animal pole stem cells. We examined if this expression signature is conserved in blastula-stage lamprey embryos. *In situ* hybridization showed that components of the neural crest (*tfap2a*, *zic-A*, *foxD-A*, *snail*, *soxE2*, *pax3/7*, *id-A*) and pluripotency GRNs (*myc*, *klf17*, *ventx/nanog*, *soxB1a*, *soxB1b*, *pou3*, *brd4*) were expressed in lamprey blastula animal pole cells, analogous to *Xenopus* (**Fig. 1c, d**; **Extended Data Fig. 1**), and double-labeling experiments further revealed extensive co-localization of neural crest and pluripotency factors in these cells (**Fig. 1e**). These results constrain the origins of the vertebrate pluripotency GRN to the last common vertebrate ancestor and highlight deeply conserved developmental and evolutionary affinities between neural crest stem cells and animal pole stem cells.

### Retention of blastula-stage GRN components in the lamprey neural crest

After finding that neural crest and pluripotency GRNs are co-expressed in lamprey animal pole cells (**Fig. 1**), we next examined if they are retained into the neural crest. The canonical pluripotency factors *myc*, *ventx/nanog*, and *klf17* were all expressed in the lamprey blastula, gastrula ectoderm, neural plate border, and neural crest, similar to their expression in *Xenopus* (**Fig. 2a**; **Extended Data Fig. 2**). Orthologs of *soxB1* were also expressed in the blastula, gastrula ectoderm and neural plate border, but then downregulated in premigratory neural crest (**Fig. 2a**; **Extended Data Fig. 2**), concomitant with a switch to *soxE* factor expression, a feature conserved with *Xenopus*^14^. Similarly, neural crest regulatory genes such as *tfap2a*, *zic-A*, *snail, and id-A*, after initially being expressed in animal pole cells, resolved to the neural plate border and neural crest, similar to *Xenopus* (**Fig. 2b**; **Extended Data Fig. 3**). Double-labeling experiments confirmed co-localization of neural crest and pluripotency factors in these cells as was observed in blastula cells (**Fig. 2c**). Notably, we did not observe axial-specific expression of pluripotency and neural crest factors, supporting recent work^15^ suggesting that the neural crest of ancestral vertebrates displayed similar expression the anteroposterior axis. (**Extended Data Fig. 4**).

**Fig. 2.**
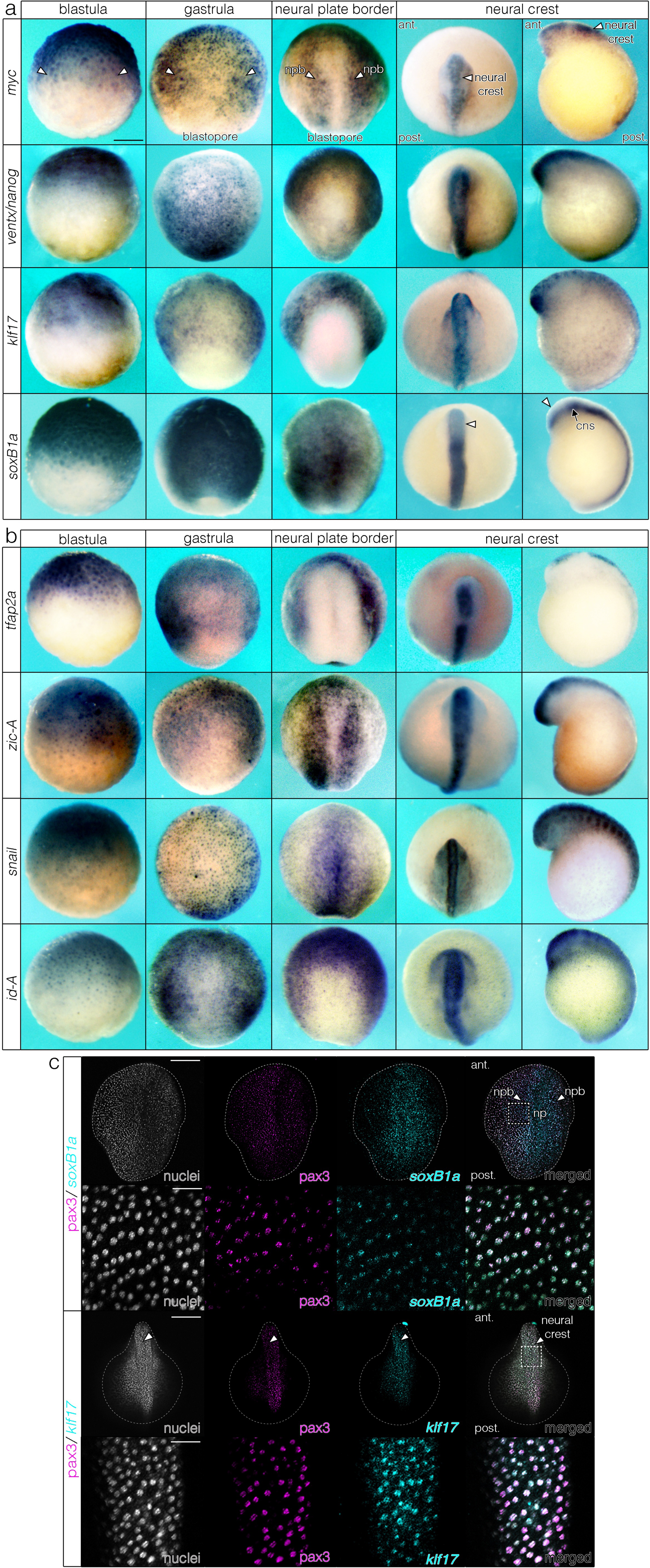
Neural crest and pluripotency GRN components are retained from the blastula to the neural plate border and neural crest in lamprey. (a, b) time series of *in situ* hybridizations for pluripotency and neural crest regulatory genes. (c) Co-localization of neural crest and pluripotency factors in the neural plate border and neural crest of lamprey. Reproducible on n ≥ 10 embryos per time point for n ≥ 3 experiments. Abbreviations: npb= neural plate border, ant = anterior, post = posterior, cns = central nervous system. Scale bar: 250 μm.

Taken together, these results show that core components of the neural crest and pluripotency GRNs of lamprey embryos are initially expressed together in animal pole cells and this expression is retained into the neural crest reminiscent to what is observed in *Xenopus*.

### Transcriptomic signature of pluripotency factors in neural crest and animal pole cells

Few studies have examined pre-gastrula stages of lamprey development^16,17^ and there are no published transcriptomes of isolated animal pole cells at these stages. The lack of such data makes it unclear if the canonical pluripotency GRN was fully present in jawless vertebrates or if some features are an innovation of jawed vertebrates. To examine this we performed RNA-Seq on animal pole explants from blastula-stage lamprey embryos (Tahara 11 (T11); **Fig. 3, Extended Data Fig. 5a-k**). Although it is unknown whether lamprey animal pole cells are functionally pluripotent (i.e., form all cell types), we recovered an expression signature consistent with such a progenitor-like state, as revealed by high levels of pluripotency factors (e.g., *myc*, *soxB1*, *klf17*, *ventx*/*nanog*) and lack of gene expression indicating commitment to germ layer differentiation (**Extended Data Fig. 5h**). We then compared our lamprey animal cap transcriptome to published transcriptomes^18^ of neural crest dissected from early (T18) and late (T21) neurula-stage embryos (**Fig. 3a**). For evolutionary comparisons, we performed RNA-Seq on pluripotent animal pole cells dissected from st9 *Xenopus* blastulae, and on animal caps reprogrammed to neural plate border (early neurula, st13) and neural crest (late neurula, st17) states using *wnt8a*/*chordin* expression (**Fig. 3a**, **Extended Data Fig. 5d-g**).

**Fig. 3.**
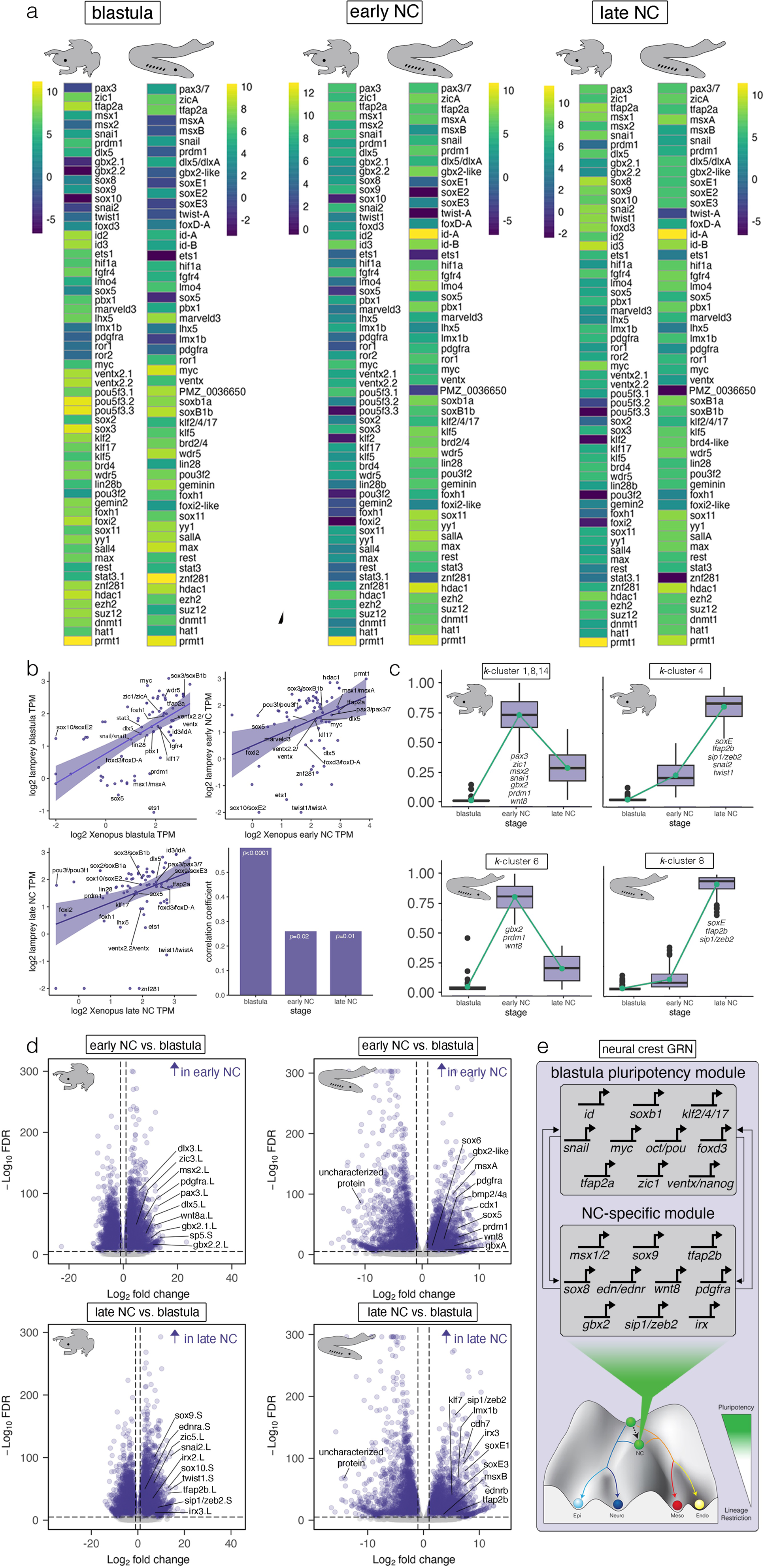
Comparative transcriptomics of neural crest and pluripotency GRNs. (a) Heatmaps of log-transformed transcript abundance (transcripts per million, TPM) for *Xenopus* and lamprey depicting expression of genes essential for pluripotency and neural crest formation. (b) Spearman correlations of TPMs across *Xenopus* and lamprey for animal pole cells and neural crest. (c) *k*-means clusters showing conserved expression dynamics of genes with an early and late neural crest signature. (d) Volcano plots showing genes that are specific to the neural crest in *Xenopus* and lamprey. Data were obtained from ≥100 lamprey animal caps (n = 3 biological replicates per stage) and ≥10 Xenopus animal caps (n = 3 biological replicates per stage). A false discovery rate of *p* < 0.05 determined statistical significance. (e) The neural crest-pluripotency GRN depicted as two distinct modules associated with blastula-stage pluripotency (top) and novel genes co-opted to the neural crest (bottom).

Directly comparing the transcriptomes of *Xenopus* and lamprey—species separated by 500 million years of independent evolution—is complicated by the lack of clear gene orthology. Therefore, gene expression levels were normalized within each species to compare the relative abundance of GRN components. We focused on neural crest factors, pluripotency factors, signaling pathways, and chromatin modifiers known to be essential for maintenance of pluripotency and/or neural crest development in vertebrates and their identified lamprey counterparts for comparisons (**Materials and Methods**). This analysis revealed considerable conservation of the relative expression levels of many of these factors, particularly within lamprey animal pole cells as compared to *Xenopus* (**Fig. 3a**). These included homologs of *id3*, *tfap2a*, *foxd3*, *pax3*, *zic1*, *lmo4*, *hif1a, dlx5, pbx1* (neural crest factors), *myc*, *stat3*, *foxh1*, *foxi2*, *soxB1 (sox2/3)*, *sox11*, *fgfr4*, *klf2/4/17*, *ventx/nanog*, *brd4*, *wdr5*, *lin28*, *znf281*, *geminin*, *yy1, sall4* (pluripotency factors), and *ezh2*, *suz12*, *hdac1*, and *prmt1* (chromatin modifiers). Although there was also evidence for conservation of relative expression levels in the neural crest, this was reduced relative to that of animal pole cells (**Fig. 3a**).

To more rigorously test evolutionary conservation, we performed cross-species correlations of transcript abundance (transcripts per million, TPM) across developmental time (**Fig. 3b**, **Materials and Methods**, **Source Data File S1**). Despite *Xenopus* and lamprey being separated by over half a billion years of independent evolution, we found statistically significant and positive cross-species correlations of transcript abundance in the blastula (*r*=0.60, *p*<0.0001), and neural crest at early neurula (*r*=0.28, *p*=0.02) and late neurula (*r*=0.29, *p*=0.01) stages (**Fig. 3b**). Notably, correlation coefficients were highest between *Xenopus* and lamprey in the blastula and lower in early and late neural crest (**Fig. 3b**), suggesting that selective constraints on transcript abundance are strongest at blastula stages. Among the most strictly conserved transcript levels across all stages were homologs of *tfap2a*, *klf17*, *soxB1*, and *ventx/nanog* (**Fig. 3b**). This suggests that strong selection pressures maintained these factors at similar levels in pluripotency GRNs across jawed and jawless vertebrates over evolutionary time. This is consistent with the requirement for precise levels of sox2 and pou5 to maintain pluripotency in mammalian ES cells^19–21^.

Our comparisons also revealed key differences in the pluripotency GRNs across vertebrates. For example, compared to *Xenopus*, lamprey *prdm1* and *sox5* show limited expression in blastula explants, and lamprey *ets1* is not expressed in these cells, as reported for the lamprey neural crest^14,15^ (**Fig. 3a**). Other important differences include moderate-to-high levels of *lefty*, *eomes*, and *otxA* in the lamprey blastula, similar to early mouse embryos^22–24^, factors which are expressed at negligible levels in *Xenopus* (**Extended Data Fig. 5i**). We also found important differences between lamprey, *Xenopus*, and mammalian pluripotency GRNs. Expression of some genes involved in mouse ES cells such as *essrb*^25,26^, *nr5a1/2*^27,28^, *cdx1/2*^29,30^, *tbx3* ^31,32^, *myb*^33^, *fgf4*^34,35^, and *prdm14* ^36,37^ was absent in lamprey and *Xenopus* blastula cells (**Extended Data Fig. 5j**). These may therefore reflect mammalian-specific adaptations of the vertebrate pluripotency GRN.

### Expression dynamics in neural crest and pluripotency GRNs

We used *k*-means clustering to compare the transcriptional dynamics of the lamprey and *Xenopus* neural crest and pluripotency GRNs. Several components showed highly correlated and conserved expression dynamics from the blastula to early and late neural crest in both species (**Fig. 3c**). For example, in both species some definitive neural crest factors, such as *soxE*, *tfap2b,* and *ednrA*/*B*, displayed monotonic increases in expression as cells progressed from blastula to late neurula stages. Interestingly, in *Xenopus* but not lamprey *twist1*, *ets1,* and *snai2* display similar expression dynamics (**Fig. 3c**). Some neural plate border factors exhibited non-monotonic expression dynamics, with their expression peaking at early neural crest stages. In both lamprey and *Xenopus*, clusters with this signature included canonical neural plate border factors such as *prdm1*, *gbx2* and *wnt8* (**Fig. 3c**). Notably, a number of other neural plate border factors, including *myc*, *pax3*, *msx1*, *zic1*, and *klf17* displayed these dynamics in *Xenopus* but not in lamprey. When we examined their expression dynamics in lamprey, we found that these genes either increased monotonically over time, peaking in late neural crest (*pax3/7*, *msx-A*, *zic-A*), or were retained at similar levels across stages (*klf17*, *snail*, *myc*) (*k*-clusters 7,8, 11, 12; **Extended Data Fig. 5l**). This suggests modifications of ancestral gene expression dynamics may have contributed to the evolution of jawed vertebrates.

### Novel neural crest GRN components co-opted into the ancestral pluripotency GRN

Beyond the regulatory signature shared with blastula stage stem cells, neural crest cells evolved in part by co-opting novel genes and signaling pathways. An example of this includes the switch from SoxB1 to SoxE factor utilization in the blastula versus neural crest, respectively^38^. To further identify such novelties shared across vertebrates, we performed differential expression analyses, focusing on transcripts that showed low expression in animal pole cells (TPMs < 10) and significant enrichment in the neural crest (**Fig. 3d, Source Data File S3**).

Our comparisons revealed overlap in differential enrichment of genes that establish and pattern the neural plate border (*wnt8*, *msx*, *gbx*, *irx*)^39–41^, initiate epithelial-mesenchymal transition (EMT) and migration (*pdgfra*, *tfap2b, sip1/zeb2, soxE*) ^42–45^, and promote lineage diversification (*endothelin* signaling and *soxE*)^46,47^ (**Fig. 3d**). We also identified species-specific differences in regulatory programs, with lamprey having neural crest-enriched expression of *prdm1*, *sox5*, and *cdh7*, whereas *Xenopus* had neural crest-enriched expression of *zic3*, *pax3*, and *dlx5* (**Fig. 3d, Extended Data Fig. 5k**). These results suggest that the neural crest evolved from the activity of at least two regulatory modules: a blastula-stage module that is also deployed in neural crest progenitors, and a neural crest-specific module co-opted downstream of the pluripotency GRN that endows the neural crest with many of its hallmark traits—EMT, migration and adoption of novel lineage states (**Fig. 3e**).

### Lamprey *pou5* is absent in neural crest but can promote neural crest formation

In jawed vertebrates, *pou5* transcription factors (pou5f1, pou5f3) are key regulators of pluripotency^48–50^. While *pou5* had been proposed to be a jawed vertebrate innovation, recently pou5 orthologs were reported in hagfish and lamprey^51–54^. Nothing is known, however, about the expression and function of *pou5* in jawless vertebrates, making it unclear if *pou5* functioned within the ancestral vertebrate pluripotency or neural crest GRNs. Our RNA-Seq analysis in lamprey revealed an uncharacterized transcript that was among the most downregulated genes in the neural crest relative to animal pole cells (**Fig. 3d**), and the top hits from a BLAST analysis were *pou5* transcription factors in jawed vertebrates. Molecular phylogenetics and synteny comparisons confirm that this gene is a lamprey ortholog of *pou5* (**Extended Data Fig. 6a, b**). In agreement with our differential expression analyses, mean transcript abundance of lamprey *pou5* was high in animal pole cells as in jawed vertebrates, but virtually absent in the neural crest (**Extended Data Fig. 6c**), an expression profile confirmed with *in situ* hybridization (**Fig. 4a**). In contrast to lamprey, the combined expression of these *Xenopus pou5f3* paralogs persists from the blastula through early neural crest stages, with maternal *pou5f3.3* expressed strongly in the blastula, and zygotic *pou5f3.1* and *pou5f3.2* expressed in the neural plate border and neural crest (**Extended Data Fig. 6d, e**).

**Fig. 4.**
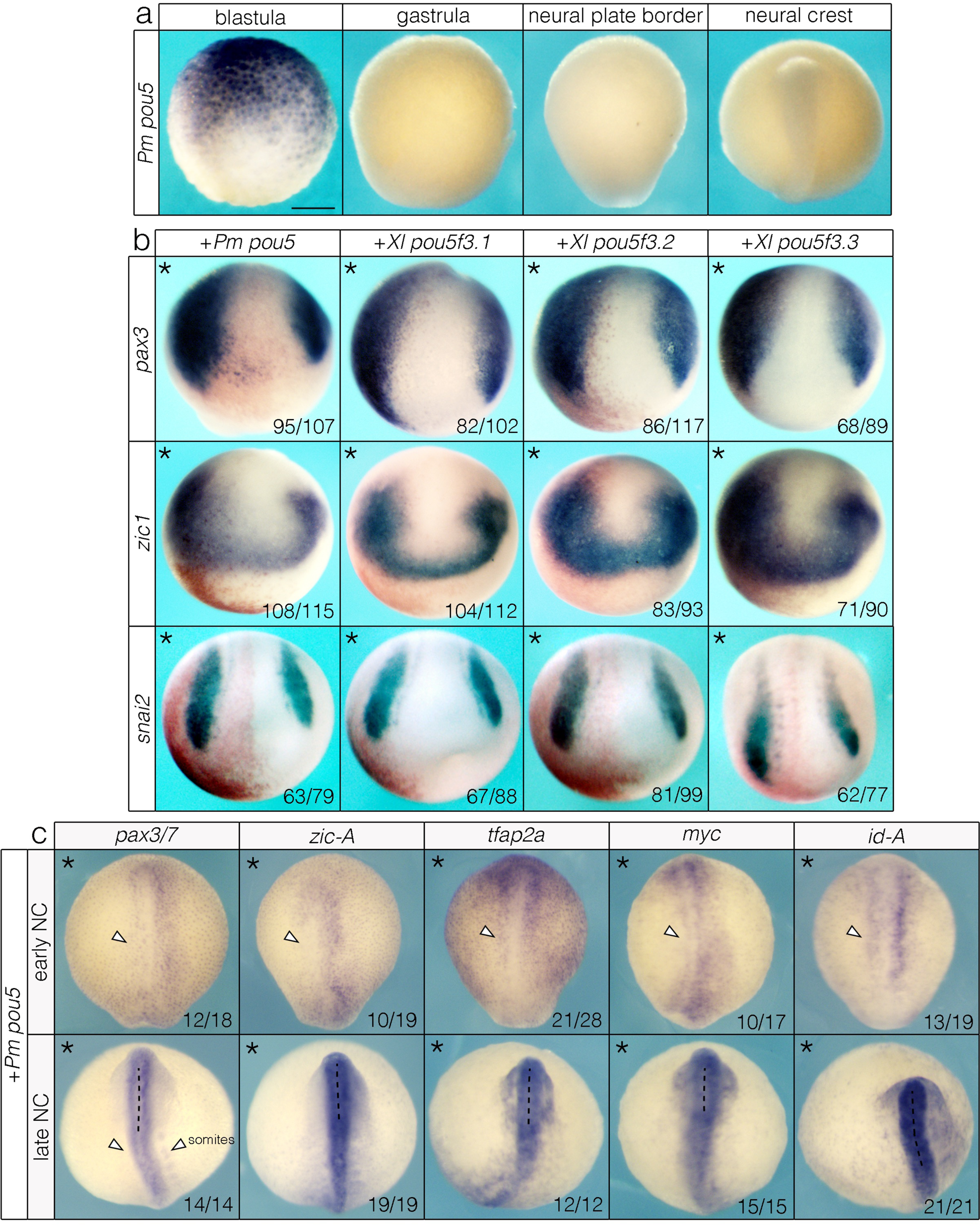
*pou5* is absent from the neural crest of jawless vertebrates but can enhance neural crest formation in a jawed vertebrate. (a) *in situ* hybridization of lamprey *pou5*. (b) Overexpression of lamprey (*Pm*) and *Xenopus* (*Xl*) *pou5* orthologs expand *pax3*, *zic1,* and *snai2* in *Xenopus* embryos. Asterisk denotes the injected side of the embryo, which is also indicated with beta-galactosidase (red) as a lineage tracer. (c) Overexpression of lamprey *pou5* in lamprey embryos either inhibits or causes no change in expression of canonical neural plate border and neural crest genes. Asterisk denotes the injected side of the embryo. Data in (b, c) were obtained from n ≥ 3 biological replicates for each construct. Scale bar: 250 μm.

Absence of *pou5* expression in the lamprey neural crest is consistent with two evolutionary scenarios: subsequent co-option into the neural crest GRN of what would become jawed vertebrates, or alternatively loss from the neural crest GRN in the lineage leading to extant jawless vertebrates. We hypothesized that if *pou5* functioned in the neural crest GRN of ancestral vertebrates but was lost in jawless vertebrates, then lamprey *pou5* may still be capable of functionally engaging the neural crest GRN. Alternatively, if lamprey *pou5* was never part of the neural crest GRN of jawless vertebrates, then expressing it in the neural crest might have neutral or antagonistic effects on neural crest formation.

To distinguish these possibilities, we expressed lamprey *pou5* mRNA on one side of early *Xenopus* embryos, comparing its activity to that of *Xenopus pou5f3.1*, *pou5f3.2*, and *pou5f3.3* expressed at equivalent levels (**Extended Data Fig. 7a**). All *pou5* factors expanded the expression of neural plate border and neural crest markers *pax3*, *zic1,* and *snai2* to comparable levels (**Fig. 4b**). Thus, the neural crest-enhancing activity of *pou5* transcription factors is an ancestral feature of *pou5.* We also reasoned that if *pou5* was secondarily lost in modern jawless vertebrates, then the lamprey neural crest GRN would have evolved independent of its activity and therefore introducing it ectopically might have neutral or antagonistic effects. To test this, we expressed lamprey *pou5* or *Xenopus pou5f3* paralogs in lamprey embryos. In contrast to *Xenopus*, we observed reduction or no change of expression of *pax3/7*, *zic-A*, *tfap2a*, *myc*, and *id-A*, (**Fig. 4c**, **Extended Data Fig. 7b**) suggesting that the ability of *pou5* to productively engage the neural crest GRN has been lost in lamprey.

### pou5 is required for neural crest formation

Because our gain-of-function experiments showed that pou5 has neural crest-enhancing activity, we next asked if pou5 is required for neural crest formation in *Xenopus*. Morpholino-(MO) mediated depletion of the zygotic *pou5f3* factors (*pou5f3.1* and *pou5f3.2*) lead to reduction of *pax3* and expansion of *zic1* at the neural plate border (**Fig. 5a**). This resulted in a near-total loss of *snai2* and *foxd3* expression (**Fig. 5a**) that was not driven by a compensatory increase in *pou5f3.3* expression (**Extended Data Fig. 8**). We then tested if lamprey *pou5* can functionally substitute for *Xenopus pou5f3* by comparing its ability to rescue the pou5 depletion phenotype compared to *Xenopus pou5f3.1* and *pou5f3.*2. We found that both lamprey and *Xenopus pou5* orthologs restored *snai2* and *foxd3* expression (**Fig. 5b**), providing further evidence that lamprey *pou5* can properly engage the neural crest GRN of jawed vertebrates.

**Fig. 5.**
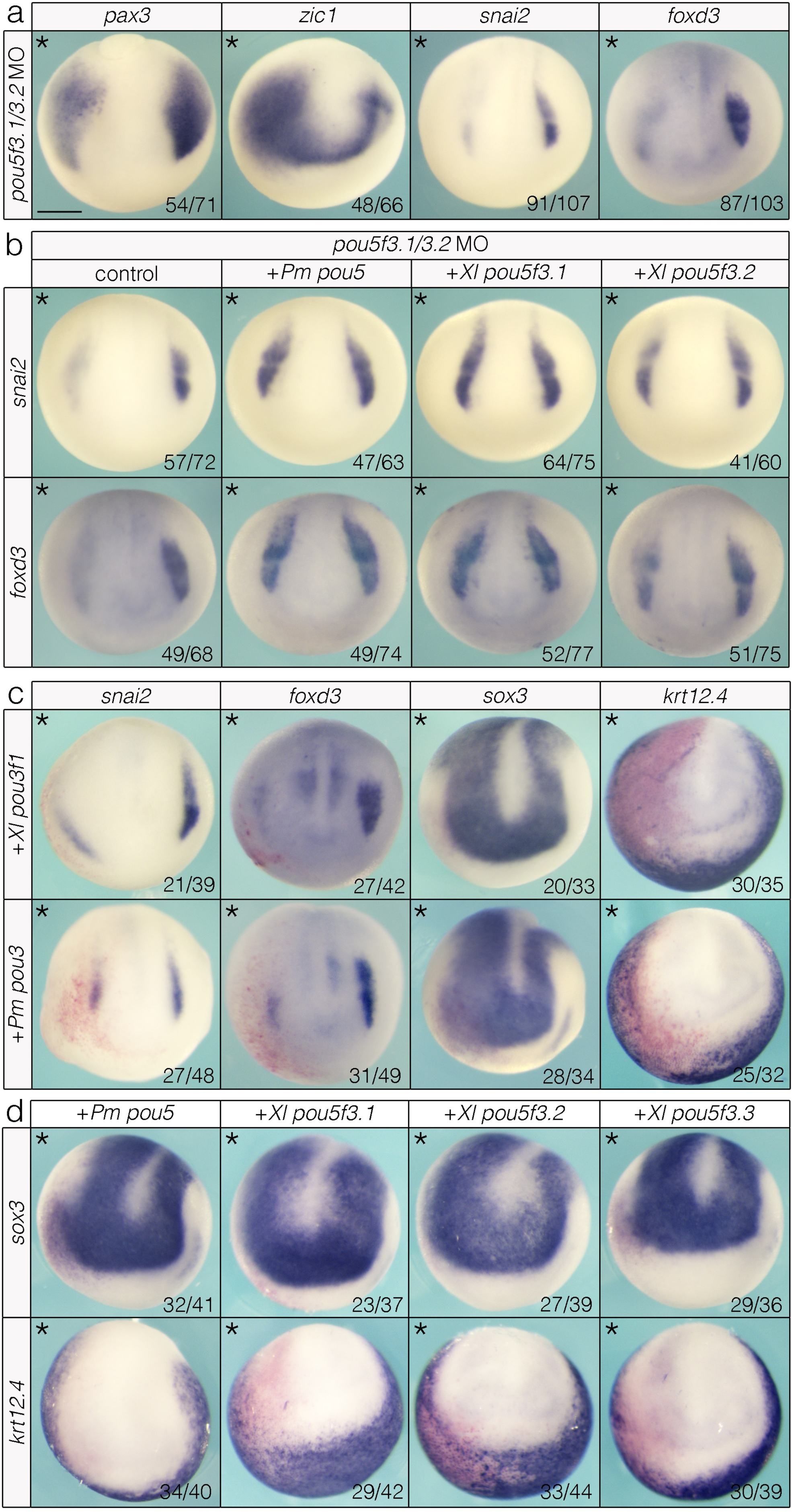
*pou5* is essential for neural crest formation and evolved neural crest-enhancing activity from a *pou3*-like ancestor involved in neurogenesis. (a) MO-mediated knockdown of zygotic *pou5f3* paralogs results in mis-patterning of the neural plate border (*pax3*, *zic1*) and loss of neural crest (*snai2*, *foxd3*). (b) lamprey and *Xenopus pou5* can rescue *pou5* MO phenotypes. (c) *pou3* gain-of-function causes loss of neural crest and epidermis and expands the neural plate. (d) *pou5* retains an ancestral role in enhancing neurogenesis. Data were obtained from n ≥ 3 biological replicates for each construct with a minimum of n = 30 embryos per replicate. Pm = *Petromyzon marinus*, Xl = *Xenopus laevis*. Scale bar: 250 μm.

### pou5 evolved neural crest-enhancing activity from a pou3-like ancestor

Based on the absence of *pou5* in invertebrate genomes, pou5 appears to be a vertebrate innovation, as with another pluripotency factor, ventx/nanog^9^. Thus, the advent of pou5, like ventx/nanog, coincides with the acquisition of neural crest and the evolution of vertebrates. However, the origins of pou5 in the neural crest remain unknown. To address this, we first revisited our phylogenetic analysis (**Extended Data Fig. 6a**), and determined that pou3 was basal to pou5, indicating that pou5 likely evolved from a pou3-like ancestor as previously suggested^52^. *pou3* has been reported to be expressed in neural progenitors, and this is the case in both lamprey and *Xenopus*, with *pou3* transcripts ultimately resolving to the neural plate (**Extended Data Fig. 9**). These results are consistent with pou5 having evolved from a pou3-like ancestor involved in neurogenesis^55,56^—a function likely ancestral for chordates and therefore one that preceded the origin of pou5 function in the neural crest.

If *pou5* evolved from a *pou3*-like ancestor, then *pou3* factors may also be capable of enhancing neural crest formation. Alternatively, the neural crest-enhancing function of *pou5* may be an evolutionary novelty. To test these possibilities, we expressed *Xenopus* and lamprey *pou3* in *Xenopus* embryos at equivalent levels. Notably, both lamprey and *Xenopus pou3* inhibited—rather than promoted—*snai2* and *foxd3* expression (**Fig. 5d**). This loss of neural crest was accompanied by a lateral expansion of the neural plate as evidenced by an expanded domain of *sox3* expression, a result consistent with pou3 factors promoting neurogenesis (**Fig. 5d**). We then asked if *pou5* factors retain an ancestral ability to promote neurogenesis. Strikingly, we found that all *pou5* orthologs expanded *sox3* expression in the neural plate (**Fig. 5e**) in addition to their neural crest-enhancing effects. By contrast these factors led to a loss of *krt12.4+* epidermal progenitors (**Fig. 5e**). Together, these results provide insights into the evolution of pou5 activity in the ancestral vertebrate neural crest GRN and point to a unique role for *pou5* in promoting neural crest development that evolved from an ancestral *pou3*-like clade involved in neurogenesis.

## Discussion

Understanding the origins of the neural crest and its developmental potential is essential to deciphering the evolution of vertebrates. Comparative approaches are critical to understanding this pivotal evolutionary transition and studies in traditional models, including *Xenopus*, chick, zebrafish, and mouse will continue to reveal species-specific differences that arose due to adaptation or developmental drift.

By comparing neural crest and pluripotency GRNs across vertebrates, we have begun to elucidate how these GRNs are developmentally and evolutionarily coupled. Our results suggest that the integration of *pou5* and *ventx/nanog* into the progenitor network of *klf17*, *soxB1,* and *myc* was already fixed in the last common vertebrate ancestor (**Fig. 6**). Although it has yet to be demonstrated that cyclostome animal pole cells are functionally pluripotent, our results indicate that these cells express the factors that control pluripotency in jawed vertebrates. Moreover, lamprey and *Xenopus* animal pole cells also express genes classically associated with neural crest identity (*tfap2a*, *snail*, *zic*, *foxd3*, *id*) supporting a model wherein multiple tiers of the neural crest GRN originated in animal pole blastula cells and were retained into the neural crest of early vertebrates. Our results also show that this neural crest-pluripotency GRN was further elaborated upon by downstream integration of novel genes and signaling pathways (e.g., *wnt8*, *msx*, *gbx*, *soxE*, *endothelins*) that pattern the neural plate border, initiate EMT, and control lineage diversification of neural crest fates. Taken together, we hypothesize that these events endowed cells at the neural plate border of stem vertebrates with the potential to layer new cell types onto the ancestral chordate baüplan, and that this pluripotency-neural crest GRN is what largely distinguishes the broad developmental potential of the vertebrate neural crest from the unipotent or bipotent cells of invertebrate chordates that have neural crest-like features^57–59^.

**Fig. 6.**
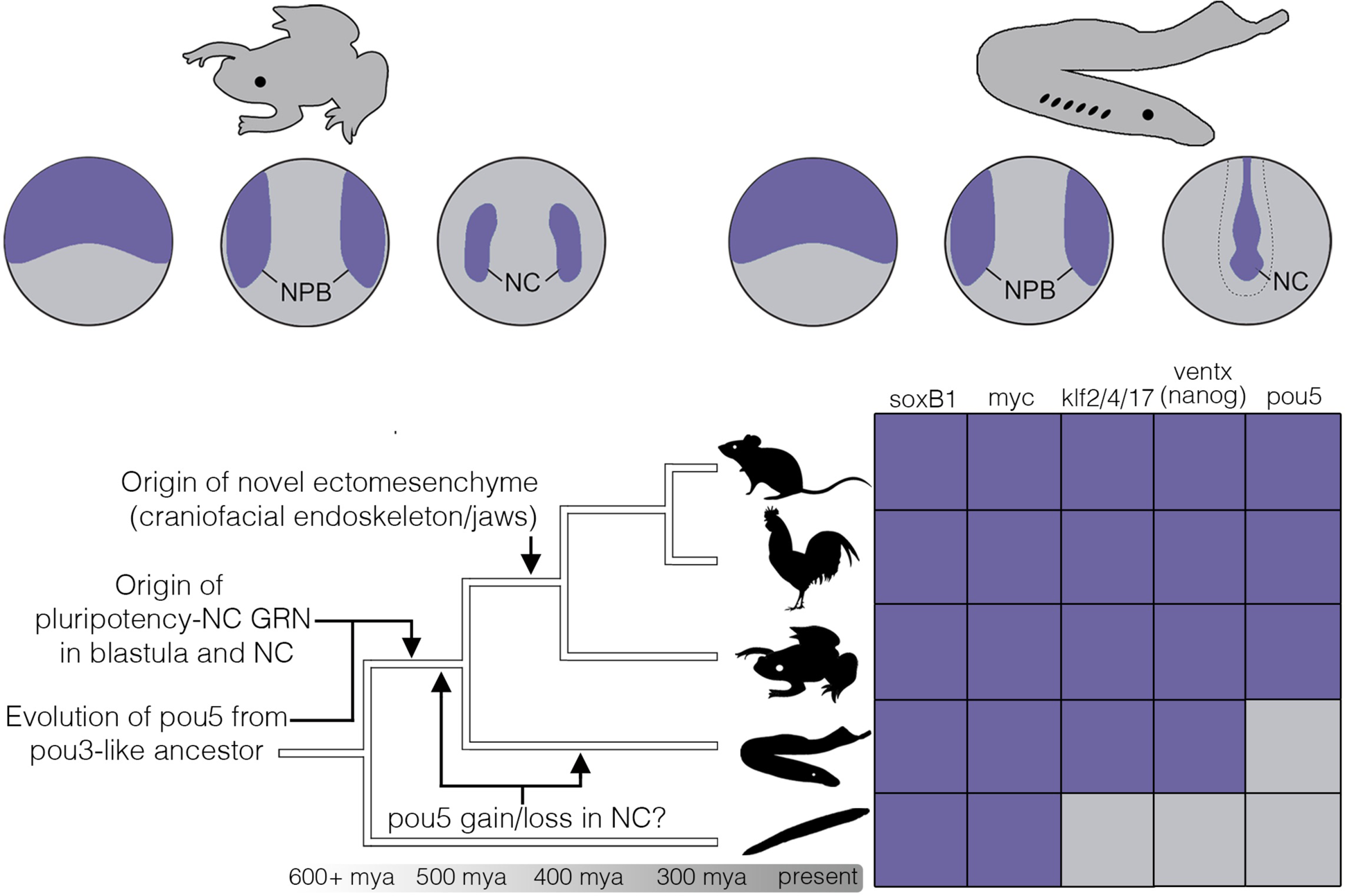
Neural crest cells evolved by retaining blastula-stage GRN components. Expression of canonical pluripotency factors in the neural crest mapped onto a chordate phylogeny. Purple boxes indicate conserved expression in the neural plate border and/or neural crest; gray boxes indicate absence of expression. NPB = neural plate border; NC = neural crest. From top to bottom, chordate lineages are represented by mouse, chick, *Xenopus*, lamprey, and invertebrate chordates.

Historically, pou5 has been recognized as a key regulator of pluripotency in ES cells, but has also been recently implicated in controlling neural crest potential^12^. Here, we show that *pou5* is not expressed in the lamprey neural crest unlike in jawed vertebrates. Lamprey neural crest also gives rise to a more limited set of derivatives and these animals lack a bony endoskeleton and jaws. It is intriguing to speculate that the broader developmental potential of the gnathostome neural crest could have been driven in part by deployment of *pou5*, highlighting yet another potential link between pluripotency factors in the neural crest and the origin of ectomesenchyme^5,9,60^. Importantly, despite being absent in the lamprey neural crest, when expressed in *Xenopus* lamprey pou5 can expand the pool of neural crest progenitors whereas it does not do so when expressed in lamprey. This suggests that *cis*-regulatory evolution of *pou5* binding sites, either through loss in cyclostomes or acquisition in jawed vertebrates, was a driver of *pou5* expression dynamics in the neural crest, coupled with rapid rates of evolution potentially driving novel protein-protein interactions^54^. Taken together with previous work showing a functional role for *pou5* in supporting blastula-stage pluripotency in *Xenopus*^61,62^, we hypothesize that pou5 activity helps maintain a similar progenitor state in the neural crest. Finally, we provide evidence that *pou5* evolved neural crest-enhancing activity from an ancestral pou3-like factor functioning in neurogenesis, and that this neural-promoting activity has been retained in both *Xenopus* and lamprey pou5 and pou3. Importantly, however, only pou5 appears to have evolved potent neural crest-promoting activity. Thus, the ability of pou5 to promote neural crest development is not a general feature of pou-family proteins but rather a synapomorphy of the pou5 clade that emerged after diverging from a pou3-like ancestor (**Fig. 6**).

Based on the recognition that the core pluripotency factors *pou5* and *ventx*/*nanog* are vertebrate novelties^9,51^ we suggest that blastula-stage (i.e., somatic) pluripotency itself is likely a vertebrate innovation. Thus, although the blastula stage is an ancient feature of metazoan development, a dual pluripotency-neural crest GRN, driven by the vertebrate-specific genes *pou5* and *ventx*/*nanog* is an evolutionary innovation of vertebrates. Evidence for this comes from the absence of the core pluripotency factors *pou5*, and *ventx*/*nanog* in invertebrate genomes, and the finding that cells of pre-blastula tunicate embryos—the sister group to vertebrates—are already restricted to a single lineage^63,64^. Although *pou5* and *ventx*/*nanog* are not encoded in invertebrate chordate genomes ^9,65,66^, invertebrate chordates do express homologs of the neural stem cell factors *soxB1* and *myc* in the blastula^17^. Thus the precursors of the vertebrate pluripotency GRN may have arisen in cells fated to be neural progenitor cells, perhaps explaining why the neural lineage has been found to lie closest in gene regulatory state space to pluripotent blastula cells and why several neural crest regulatory factors have been found to have deep roots in bilaterian neurogenesis^4,67,68^.

## Methods and Materials

### Embryological methods

Adult sea lamprey were collected from the Hammond Bay Biological Station, Millersburg, MI, and shipped to Northwestern University. Animals were maintained at 14 °C in a recirculating water system. Embryos were obtained by *in vitro* fertilization and cultured in 0.05X Marc’s Modified Ringers (MMR) in Pyrex dishes. All procedures were approved by Northwestern University’s Institutional Animal Care and Use Committee (IACUC A3283-01). Blastula-stage animal pole explants were manually dissected from lamprey embryos that were dechorionated in 1X MMR in a dish lined with 1% agarose. Animal caps were dissected manually using sharp forceps and cultured until the desired stage before harvesting for total RNA extraction. For *Xenopus*, controls were wildtype (epidermal) caps collected at st 13 and st 17. Alternatively, caps were induced to form neural plate border (st 13) and neural crest (st 17) by microinjection of *wnt8a* and *chordin* mRNA at the 2-cell stage. mRNA for microinjection was *in vitro* transcribed from a linearized DNA template using the SP6 mMessage mMachine kit (ThermoFisher AM1344). All animal cap experiments were collected for three biological replicates.

### Cloning and *in situ* hybridization

*X. laevis* clones were obtained from ORFeome (www.xenbase.org/reagents/static/orfeome.jsp). Lamprey clones were obtained from previous library screens or gene synthesis (Synbio Technologies). *in situ* hybridization was performed as done previously (*11,12*). Embryos and sections were imaged on an Olympus SZX12 microscope equipped with an Olympus QColor3 camera and QCapture software.

### Double labeling with HCR-FISH with immunohistochemistry

For HCR-FISH, we adopted the third generation HCRv3-FISH^69^ protocol. HCR-FISH probe sets targeting were custom-designed by Molecular Instruments. Immediately after HCR-FISH, lamprey embryos were rinsed in PBS-T (1x PBS, 0.1% Tween 20), blocked (PBS-T, 10% fetal bovine serum), and then rocked overnight at 4°C in anti-Pax3 primary antibody that has been previously validated in lamprey^70^ (DSHB #528426, 1:5). After several rinses in PBS-T, samples were incubated in goat anti-mouse IgG Alexa Fluor™ 546 secondary antibody (ThermoFisher A28182), rinsed again several times in PBS-T, and incubated with SYTOX Green nucleic acid stain (ThermoFisher S7020). Samples were mounted and imaged using a Nikon C2 confocal microscope. Colocalization was determined by examining individual optical sections of z-slices through a single cell depth.

### RNA isolation, library preparation, and sequencing

Total RNA from *Xenopus* and lamprey was extracted using TRIZOL reagent (ThermoFisher 15596026) and LiCl precipitation as per standard procedures. Lamprey RNA-Seq libraries were prepared using the NEBNext Ultra II Directional RNA Library Prep Kit for Illumina, along with the NEBNext Poly(A) mRNA Magnetic Isolation Module and NEBNext High-Fidelity 2X PCR Master Mix using the manufacturer’s protocol. *Xenopus* RNA-Seq libraries were prepared using the Illumina TruSeq Library Prep kit. Libraries were quantified by Qubit and assessed using an Agilent TapeStation. Next Generation Sequencing was performed at the NUSeq Core facility at Northwestern Medicine’s Center for Genetic Medicine. We obtained 50 million single end reads on three biological replicates for lamprey animal caps and *Xenopus* animal cap experiments on the Illumina NextSeq500 platform. For previously published lamprey neural crest RNA-Seq data sets (T18 neural folds, T20, T21 dorsal neural tubes), we downloaded FASTQ files from the European Nucleotide Archive (https://www.ebi.ac.uk/ena/browser/home) and processed these files in parallel with our lamprey animal cap transcriptomes as outlined below.

### Processing and analysis of RNA-Seq data

RNA-Seq read quality was evaluated using FASTQC (v0.11.5). Reads were trimmed by fastp (v0.23.2) for quality and to remove Illumina adapters using default parameters. Trimmed reads were mapped to the *Xenopus* (Xl9.2) (https://www.xenbase.org/other/static-xenbase/ftpDatafiles.jsp) or sea lamprey germline genome assemblies (https://simrbase.stowers.org/sealamprey) using STAR (v2.6.0). Read counts were obtained using HTSeq (v1.99.2). PCA and differential expression analyses were performed in DESeq2 (v3.14) using R (v4.0.3; http://www.R-project.org/). For comparisons of transcriptomes between *Xenopus* and lamprey, we constructed heatmaps (‘pheatmap’ package in R) using log-transformed normalized transcript abundance values (transcripts per million, TPM) generated from RSEM (v1.2.28). This method is ideal for our comparisons as it allows for normalized transcript abundance of genes to be directly compared across species. For our comparisons, we curated from the literature a set of transcription factors, signaling pathways, and epigenetic modifiers known to be essential for pluripotency and/or neural crest development in vertebrates.

To test evolutionary conservation of transcript abundance between *Xenopus* and lamprey, we performed correlation analysis on the transcripts in in **Fig. 3a**. For genes having a single putative ortholog in both species (e.g., *Xenopus foxd3* and lamprey *foxD-A*), we calculated the mean TPMs for each gene in each cell population for one-to-one comparisons. For paralogous groups of genes where one-to-one orthology is unclear or unknown, we performed pairwise comparisons between genes within each paralogy group. For example, the lamprey genome encodes a single *snail* ortholog (*11*), which has affinities to both *snai1* and *snai2*, whereas *Xenopus* has distinct *snai1* and *snai2* paralogs. In such cases, for example, the mean TPMs for lamprey *snail* in each cell population was compared separately to both *snai1* and *snai2* in *Xenopus*. We performed separate correlation analyses (blastula, early neural crest, late neural crest) between *Xenopus* and lamprey, using log transformed TPMs. Visual inspection of residual plots from parametric correlation analyses revealed significant departures from normality. We therefore performed non-parametric Spearman rank correlation analyses in R (v4.0.3) using the ‘cor.test’ function (‘stats’ package). We report correlation coefficients from our analyses; effects were statistically significant where *p*<0.05. For *k*-means analysis, RNA-Seq data were analyzed using the package ‘coseq’ with the following parameters: K=2:25, transformation=“logclr”,norm=“DESeq”,meanFilterCutoff=50, model=“kmeans”, iter.max=10,seed=12345.

To test for potential batch effects from incorporating our lamprey RNA-Seq data with previously published data sets (described above), we performed PCA with two additional T21 dorsal neural tube replicates from another independent publication^15^. We performed the same analyses in *Xenopus* by incorporating an additional neurula-stage samples in which neural crest was dissected from the embryo ^71^. The results from these analyses indicate that differences are primarily biological and lack significant batch effects (**Extended Data Fig. S5**).

### Phylogenetic analysis and synteny of vertebrate pou5 transcription factors

Full length oct/pou-family proteins were downloaded manually from NCBI. Sequences were aligned using MAFFT (v7.490) with <--maxiterate 1000 –-globalpair>, and then trimmed using trimAl (v1.4.1). The <-automated1> option was used to heuristically determine the optimal method for trimming. The trimmed alignment file was converted to NEXUS format for phylogenetic analysis in MrBayes (v3.2.7a), or PHYLIP format for analysis in RAxML (v8). The following parameters were used in MrBayes: <prset aamodel = mixed>, <mcmc ngen = 500000>; mouse pou6f1 was specified as outgroup. The following parameters were used in RAxML: <-m PROTGAMMAAUTO>, <-p12345>, <-x 12345>, <-#1000>; mouse pou6f1 was specified as outgroup. Consensus trees were visualized using iTOL (https://itol.embl.de). NCBI accession numbers for oct/pou sequences are: *Ambystoma mexicanum* pou5f3 AGN30963.1; *Ambystoma mexicanum* pou5f1 AY542376.1; *Danio rerio* pou1f1 NP_998016.1; *Danio rerio* pou5f3 BAA05901.1; *Danio rerio* pou2f2 XP_009290402.1; *Danio rerio* pou2f3 XP_005172050.1; *Danio rerio* pou3f1 NP_571236.1; *Danio rerio* pou3f2 NP_571235.1; *Danio rerio* pou3f3 NP_571225.2; *Danio rerio* pou4f1 NP_001299795.1; *Felis catus* pou5f1 ACY72350.1; *Gallus gallus* pou5f3 ABK27428.1; *Gallus gallus* pou2f1 NP_990803.1; *Gallus gallus* pou2f3 XP_015153671.1; *Gallus gallus* pou3f1 NP_001026755.1; *Gallus gallus* pou3f2 XP_015140157.1; *Gallus gallus* pou3f3 XP_040518115.1; *Gallus gallus* pou3f4 XP_003641125.3; *Gallus gallus* pou4f1 XP_015132813.1; *Homo sapiens* pou5f1 BAC54946.1; *Mus musculus* pou1f1 AAH61213.1; *Mus musculus* pou2f1 NP_001355737.1; *Mus musculus* pou2f2 NP_001157028.1; *Mus musculus* pou2f3 NP_035269.2; *Mus musculus* pou3f1 NP_035271.1; *Mus musculus* pou3f2 NP_032925.1; *Mus musculus* pou3f3 NP_032926.2; *Mus musculus* pou3f4 NP_032927.1; *Mus musculus* pou4f1 NP_035273.3; *Mus musculus* pou4f2 NP_620394.2; *Mus musculus* pou5f1 NP_038661.2; *Oryzias latipes* pou5f3 AAT64911.1; *Petromyzon marinus* pou5 XP_032835430.1; *Petromyzon marinus* pou3f2-like XP_032828582.1; *Petromyzon marinus* pou2f2-like XP_032821400.1; *Petromyzon marinus* pou2f1 XP_032833038.1; *Petromyzon marinus* pou2f2 XP_032821327.1; *Xenopus laevis* pou5f3.1 NP_001081342.1; *Xenopus laevis* pou5f3.2 NP_001079832.1; *Xenopus laevis* pou5f3.3 NP_001081583.1; *Xenopus tropicalis* pou1f1 XP_031752450.1; *Xenopus tropicalis* pou2f1 XP_012812406.1; *Xenopus tropicalis* pou2f2 XP_002936445.2; *Xenopus tropicalis* pou2f3 XP_031762183.1; *Xenopus tropicalis* pou3f1 NP_001016504.1; *Xenopus tropicalis* pou3f2 NP_001263306.1; *Xenopus tropicalis* pou3f3 XP_017947125.1; *Xenopus tropicalis* pou3f4 NP_001090728.1; *Xenopus tropicalis* pou4f1 XP_002931972.1; *Mus musculus* pou6f1 AAH85139.1. Other sequences were obtained from previously published data sets^54^. Synteny analysis was performed by comparing the coding sequences of *Xenopus* genes surrounding the *pou5f3.1*, *pou5f3.2*, and *pou5f3.3* loci to those surrounding the *pou5* locus in the germline genome assembly of lamprey.

### Overexpression of lamprey and *Xenopus* pou5 and pou3 mRNA in *Xenopus* embryos

Full-length lamprey *pou5*, lamprey *pou3*, *Xenopus pou5f3.1*, *Xenopus pou5f3.2*, *Xenopus pou5f3.3*, and *Xenopus pou3f1* were subcloned into the pCS2 vector with a 6X N-terminal MYC tag. mRNA for microinjection was *in vitro* transcribed from linearized DNA templates using an SP6 mMessage mMachine SP6 kit. Injections were performed at the two- or four-cell stage with beta-galactosidase as a lineage tracer. Protein levels were matched for phenotypic comparisons as determined by Western analysis (**Extended Data Fig. 8**) with an anti-MYC antibody as described in (*18*).

### pou5 morpholino experiments in *Xenopus*

Previously validated FITC-conjugated morpholino (MO) oligonucleotides^61,62^ targeting *Xenopus pou5f3.1* and *pou5f3.2* were injected unilaterally (∼1.4 ng per MO) into one or two animal pole blastomeres at the eight-cell stage with or without MYC-tagged mRNA encoding full-length lamprey *pou5*, *Xenopus pou5f3.1*, or *pou5f3.2*. Embryos were sorted for unilateral incorporation of FITC. MO sequences: *pou5f3.1.L* = CCTATACAGCTCTTGCTCAAATC, *pou5f3.1.S* = GATTAAACATGATCTGTTGTCCG, *pou5f3.2.L* = CCAAGAGCTTGCAGTCAGATC, *pou5f3.2.S* = GCTGAACCCTAGAATGACCAG

### Overexpression of lamprey pou5 mRNA in lamprey embryos

Lamprey embryos were injected with MYC-tagged lamprey (*pou5*) or *Xenopus* (*pou5f3.1*, *pou5f3.2*, *pou5f3.3*) mRNA and a FITC dextran tracer (ThermoFisher AAJ6360622) into one blastomere at the two-cell stage at the same concentration used for injections in *Xenopus*. In approximately 30-40% of lamprey embryos, the first cleavage prefigures the left-right axis. Therefore, embryos were sorted by unilateral FITC incorporation before processing for *in situ* hybridization.

### Sectioning

Lamprey embryos that were processed by *in situ* hybridization were embedded in 4% agarose, and then sectioned on a Leica VT100S vibratome.

## Acknowledgments

We thank Scott Miehls and the staff of the Hammond Bay Biological Station for shipment of lampreys, David McCauley, Daniel Medeiros, Tatjana Sauka-Spengler, and Marianne Bronner for clones and reagents, and Rosemary Braun for statistical advice. We also thank participants in the Woods Hole Embryology course and members of the LaBonne laboratory for helpful discussions.

## Author contributions

Conceptualization: JRY, CL

Methodology: JRY, CL

Investigation: JRY, AR, PH, AM, ENS, SR

Visualization: JRY

Funding acquisition: CL, JRY

Project administration: JRY, CL

Supervision: CL

Writing – original draft: JRY

Writing – review & editing: JRY, AR, PH, AM, ENS, CL, SR

## Data and materials availability

RNA-Seq data have been deposited at NCBI (GSE205436) All other data are available in the main text or the supplementary materials.

## Extended Data

Extended Data Figs. S1 to S9

Extended Data Source Files S1, S2

## References

1 Buitrago-Delgado, E., Nordin, K., Rao, A., Geary, L. & LaBonne, C. Shared regulatory programs suggest retention of blastula-stage potential in neural crest cells. Science 348, 1332–1335 (2015).

2 Le Douarin, N. & Kalcheim, C. The neural crest. 2nd edn, (Cambridge ; New York : Cambridge University Press, 1999).

3 Schock, E. N., York, J. R. & LaBonne, C. in Semin. Cell Dev. Biol. (Elsevier).

4 York, J. R. & McCauley, D. W. The origin and evolution of neural crest cells. Open Biology 10, 190285 (2020).

5 Green, S. A., Simões-Costa, M. & Bronner, M. Evolution of vertebrates as viewed from the crest. Nature 520, 474–482, doi:10.1038/nature14436 (2015).

6 Lignell, A., Kerosuo, L., Streichan, S. J., Cai, L. & Bronner, M. E. Identification of a neural crest stem cell niche by Spatial Genomic Analysis. Nature Communications 8, 1–11 (2017).

7 Nordin, K. & Labonne, C. Sox5 Is a DNA-Binding Cofactor for BMP R-Smads that Directs Target Specificity during Patterning of the Early Ectoderm. Developmental Cell 31, 374–382, doi:10.1016/j.devcel.2014.10.003 (2014).

8 Scerbo, P. et al. Ventx factors function as Nanog-like guardians of developmental potential in Xenopus. PLoS One 7, e36855 (2012).

9 Scerbo, P. & Monsoro-Burq, A. H. The vertebrate-specific VENTX/NANOG gene empowers neural crest with ectomesenchyme potential. Science Advances 6, eaaz1469 (2020).

10 Rao, A. & LaBonne, C. Histone deacetylase activity has an essential role in establishing and maintaining the vertebrate neural crest. Development 145, dev163386 (2018).

11 Geary, L. & LaBonne, C. FGF mediated MAPK and PI3K/Akt Signals make distinct contributions to pluripotency and the establishment of Neural Crest. Elife 7, e33845 (2018).

12 Zalc, A. et al. Reactivation of the pluripotency program precedes formation of the cranial neural crest. Science 371, eabb4776 (2021).

13 Pajanoja, C. et al. Maintenance of pluripotency-like signature in the entire ectoderm leads to neural crest stem cell potential. Nature Communications (2023).

14 Sauka-Spengler, T., Meulemans, D. M., Jones, M. & Bronner-Fraser, M. Ancient evolutionary origin of the neural crest gene regulatory network. Developmental Cell 13, 405–420, doi:10.1016/j.devcel.2007.08.005 (2007).

15 Martik, M. L. et al. Evolution of the new head by gradual acquisition of neural crest regulatory circuits. Nature 574, 675–678 (2019).

16 Takeuchi, M., Takahashi, M., Okabe, M. & Aizawa, S. Germ layer patterning in bichir and lamprey: an insight into its evolution in vertebrates. Developmental Biology 332, 90–102 (2009).

17 Cattell, M. V., Garnett, A. T., Klymkowsky, M. W. & Medeiros, D. M. A maternally established SoxB1/SoxF axis is a conserved feature of chordate germ layer patterning. Evol. Dev. 14, 104–115 (2012).

18 Hockman, D. et al. A genome-wide assessment of the ancestral neural crest gene regulatory network. Nature Communications 10, 1–15 (2019).

19 Shi, G. & Jin, Y. Role of Oct4 in maintaining and regaining stem cell pluripotency. Stem cell research & therapy 1, 1–9 (2010).

20 Thomson, M. et al. Pluripotency factors in embryonic stem cells regulate differentiation into germ layers. Cell 145, 875–889 (2011).

21 Radzisheuskaya, A. et al. A defined Oct4 level governs cell state transitions of pluripotency entry and differentiation into all embryonic lineages. Nature cell biology 15, 579–590 (2013).

22 Kim, D.-K., Cha, Y., Ahn, H.-J., Kim, G. & Park, K.-S. Lefty1 and lefty2 control the balance between self-renewal and pluripotent differentiation of mouse embryonic stem cells. Stem Cells Dev. 23, 457–466 (2014).

23 Tosic, J. et al. Eomes and Brachyury control pluripotency exit and germ-layer segregation by changing the chromatin state. Nature Cell Biology 21, 1518–1531 (2019).

24 Acampora, D., Di Giovannantonio, L. G. & Simeone, A. Otx2 is an intrinsic determinant of the embryonic stem cell state and is required for transition to a stable epiblast stem cell condition. Development 140, 43–55 (2013).

25 Ivanova, N. et al. Dissecting self-renewal in stem cells with RNA interference. Nature 442, 533–538 (2006).

26 Zhang, X., Zhang, J., Wang, T., Esteban, M. A. & Pei, D. Esrrb activates Oct4 transcription and sustains self-renewal and pluripotency in embryonic stem cells. Journal of Biological Chemistry 283, 35825–35833 (2008).

27 Guo, G. & Smith, A. A genome-wide screen in EpiSCs identifies Nr5a nuclear receptors as potent inducers of ground state pluripotency. Development 137, 3185–3192 (2010).

28 Gassler, J. et al. Zygotic genome activation by the totipotency pioneer factor Nr5a2. Science, eabn7478 (2022).

29 Blij, S., Parenti, A., Tabatabai-Yazdi, N. & Ralston, A. Cdx2 efficiently induces trophoblast stem-like cells in naïve, but not primed, pluripotent stem cells. Stem Cells Dev. 24, 1352–1365 (2015).

30 Rousso, S. Z. et al. Negative autoregulation of Oct3/4 through Cdx1 promotes the onset of gastrulation. Developmental Dynamics 240, 796–807 (2011).

31 Han, J. et al. Tbx3 improves the germ-line competency of induced pluripotent stem cells. Nature 463, 1096–1100 (2010).

32 Russell, R. et al. A dynamic role of TBX3 in the pluripotency circuitry. Stem Cell Reports 5, 1155–1170 (2015).

33 Tanaka, Y., Patestos, N. P., Maekawa, T. & Ishii, S. B-myb is required for inner cell mass formation at an early stage of development. Journal of Biological Chemistry 274, 28067–28070 (1999).

34 Fernandez-Tresguerres, B. et al. Evolution of the mammalian embryonic pluripotency gene regulatory network. Proceedings of the National Academy of Sciences 107, 19955–19960 (2010).

35 Tanaka, S., Kunath, T., Hadjantonakis, A.-K., Nagy, A. & Rossant, J. Promotion of trophoblast stem cell proliferation by FGF4. Science 282, 2072–2075 (1998).

36 Yamaji, M. et al. PRDM14 ensures naive pluripotency through dual regulation of signaling and epigenetic pathways in mouse embryonic stem cells. Cell Stem Cell 12, 368–382 (2013).

37 Grabole, N. et al. Prdm14 promotes germline fate and naive pluripotency by repressing FGF signalling and DNA methylation. EMBO Reports 14, 629–637 (2013).

38 Buitrago-Delgado, E., Schock, E. N., Nordin, K. & LaBonne, C. A transition from SoxB1 to SoxE transcription factors is essential for progression from pluripotent blastula cells to neural crest cells. Developmental Biology 444, 50–61 (2018).

39 Monsoro-Burq, A.-H., Wang, E. & Harland, R. Msx1 and Pax3 Cooperate to Mediate FGF8 and WNT Signals during Xenopus Neural Crest Induction. Developmental Cell 8, 167–178, doi:10.1016/j.devcel.2004.12.017 (2005).

40 Li, B., Kuriyama, S., Moreno, M. & Mayor, R. The posteriorizing gene Gbx2 is a direct target of Wnt signalling and the earliest factor in neural crest induction. Development 136, 3267–3278 (2009).

41 LaBonne, C. & Bronner-Fraser, M. Neural crest induction in Xenopus: evidence for a two-signal model. Development 125, 2403–2414 (1998).

42 Lander, R. et al. Interactions between Twist and other core epithelial–mesenchymal transition factors are controlled by GSK3-mediated phosphorylation. Nature communications 4, 1–11 (2013).

43 Mancilla, A. & Mayor, R. Neural crest formation in *Xenopus laevis*: mechanisms of Xslug induction. Developmental Biology 177, 580–589 (1996).

44 Rogers, C. D., Saxena, A. & Bronner, M. E. Sip1 mediates an E-cadherin-to-N-cadherin switch during cranial neural crest EMT. Journal of Cell Biology 203, 835–847 (2013).

45 LaBonne, C. & Bronner-Fraser, M. Snail-related transcriptional repressors are required in Xenopus for both the induction of the neural crest and its subsequent migration. Developmental Biology 221, 195–205 (2000).

46 Square, T. A. et al. Evolution of the endothelin pathway drove neural crest cell diversification. Nature 585, 563–568 (2020).

47 Haldin, C. E. & LaBonne, C. SoxE factors as multifunctional neural crest regulatory factors. The international journal of biochemistry & cell biology 42, 441–444 (2010).

48 Tapia, N. et al. Reprogramming to pluripotency is an ancient trait of vertebrate Oct4 and Pou2 proteins. Nature communications 3, 1–12 (2012).

49 Nichols, J. et al. Formation of pluripotent stem cells in the mammalian embryo depends on the POU transcription factor Oct4. Cell 95, 379–391 (1998).

50 Niwa, H., Miyazaki, J.-i. & Smith, A. G. Quantitative expression of Oct-3/4 defines differentiation, dedifferentiation or self-renewal of ES cells. Nature genetics 24, 372–376 (2000).

51 Onichtchouk, D. Evolution and functions of Oct4 homologs in non-mammalian vertebrates. Biochimica et Biophysica Acta (BBA)-Gene Regulatory Mechanisms 1859, 770–779 (2016).

52 Gold, D. A., Gates, R. D. & Jacobs, D. K. The early expansion and evolutionary dynamics of POU class genes. Molecular Biology and Evolution 31, 3136–3147 (2014).

53 Bakhmet, E. I. & Tomilin, A. N. The functional diversity of the POUV-class proteins across vertebrates. Open Biology 12, 220065 (2022).

54 Sukparangsi, W. et al. Evolutionary Origin of Vertebrate OCT4/POU5 Functions in Supporting Pluripotency. Nature Communications 5537 (2022) (2022).

55 Zhu, Q. et al. The transcription factor Pou3f1 promotes neural fate commitment via activation of neural lineage genes and inhibition of external signaling pathways. Elife 3, e02224 (2014).

56 Cosse-Etchepare, C. et al. Pou3f transcription factor expression during embryonic development highlights distinct pou3f3 and pou3f4 localization in the Xenopus laevis kidney. International Journal of Developmental Biology 62, 325–333 (2018).

57 Stolfi, A., Ryan, K., Meinertzhagen, I. A. & Christiaen, L. Migratory neuronal progenitors arise from the neural plate borders in tunicates. Nature 527, 371, doi:10.1038/nature15758 (2015).

58 Abitua, P. B., Wagner, E., Navarrete, I. A. & Levine, M. Identification of a rudimentary neural crest in a non-vertebrate chordate. Nature 492, 104–107, doi:10.1038/nature11589 (2012).

59 Jeffery, W. R., Strickler, A. G. & Yamamoto, Y. Migratory neural crest-like cells form body pigmentation in a urochordate embryo. Nature 431, 696–699 (2004).

60 Smith, J. J. et al. The sea lamprey germline genome provides insights into programmed genome rearrangement and vertebrate evolution. Nature Genetics 50, 270–277 (2018).

61 Morrison, G. M. & Brickman, J. M. Conserved roles for Oct4 homologues in maintaining multipotency during early vertebrate development. Development (2006).

62 Livigni, A. et al. A conserved Oct4/POUV-dependent network links adhesion and migration to progenitor maintenance. Current Biology 23, 2233–2244 (2013).

63 Satou, Y. A gene regulatory network for cell fate specification in Ciona embryos. Current Topics in Developmental Biology 139, 1–33 (2020).

64 Lemaire, P. Unfolding a chordate developmental program, one cell at a time: invariant cell lineages, short-range inductions and evolutionary plasticity in ascidians. Developmental Biology 332, 48–60 (2009).

65 Frankenberg, S. & Renfree, M. B. On the origin of POU5F1. BMC biology 11, 1–12 (2013).

66 Vanni, V. et al. Yamanaka Factors in the budding tunicate Botryllus schlosseri show a shared spatio-temporal expression pattern in chordates. Frontiers in Cell and Developmental Biology, 394.

67 Johnson, K., Freedman, S., Braun, R. & LaBonne, C. Quantitative analysis of transcriptome dynamics provides novel insights into developmental state transitions. BMC genomics 23, 723 (2022).

68 York, J. R., Zehnder, K., Yuan, T., Lakiza, O. & McCauley, D. W. Evolution of Snail-mediated regulation of neural crest and placodes from an ancient role in bilaterian neurogenesis. Developmental Biology 453, 180–190 (2019).

69 Choi, H. M. et al. Third-generation in situ hybridization chain reaction: multiplexed, quantitative, sensitive, versatile, robust. Development 145, dev165753 (2018).

70 Migocka-Patrzałek, M. et al. Unique Features of River Lamprey (Lampetra fluviatilis) Myogenesis. International Journal of Molecular Sciences 23, 8595 (2022).

