## Supplemental data for "Shared features of blastula and neural crest stem cells evolved at the base of vertebrates"

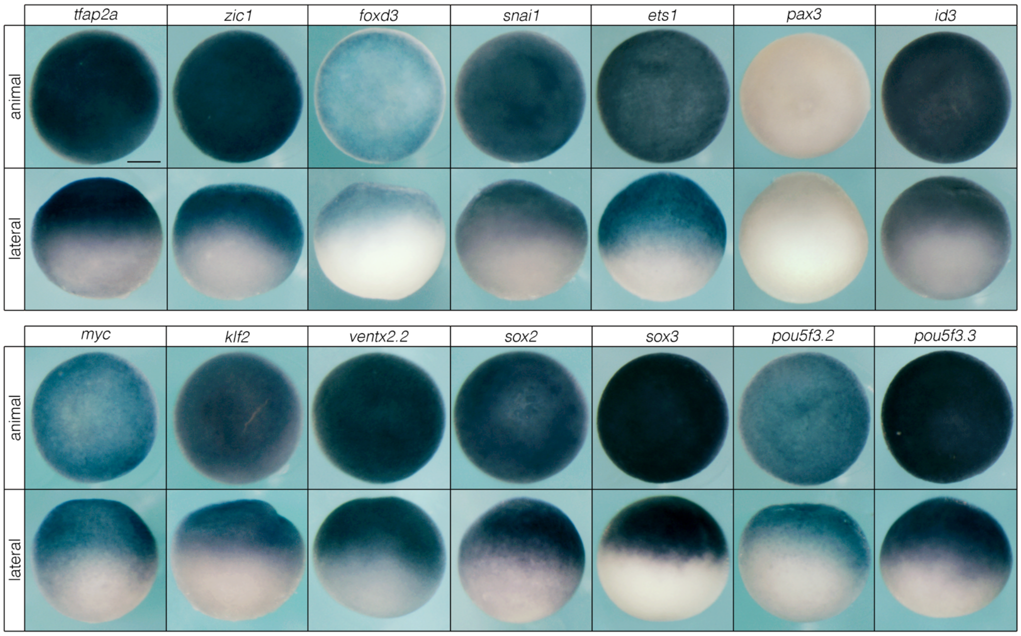


**Extended Data Fig. S1**. Neural crest and pluripotency factors are expressed in animal pole blastula cells in *Xenopus*. *In situ* hybridizations of neural crest (top) and pluripotency (bottom) factors in animal pole stem cells. Reproducible on n ≥ 10 embryos for n ≥ 3 experiments.

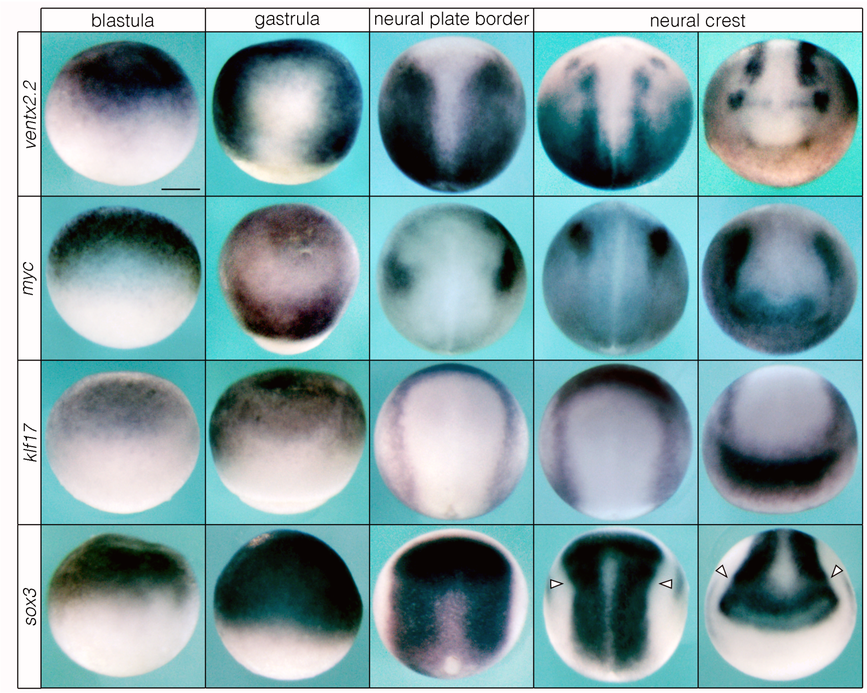


**Extended Data Fig. S2**. Pluripotency factors in the blastula gradually resolve to the neural plate border and neural crest in *Xenopus*. Arrowheads denote expression of *sox3* largely excluded from the neural crest at late neurula stages. Reproducible on n ≥ 10 embryos per time point for n ≥ 3 experiments. Scale bar: 250 µm.

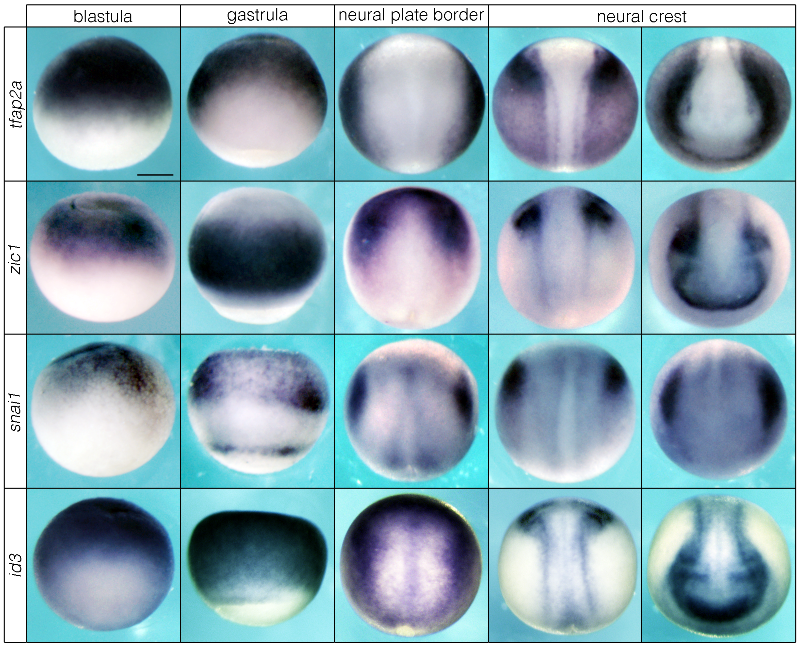


**Extended Data Fig. S3**. Neural crest factors in the blastula gradually resolve to the neural plate border and neural crest in *Xenopus*. Reproducible on n ≥ 10 embryos per time point for n ≥ 3 experiments. Scale bar: 250 µm.

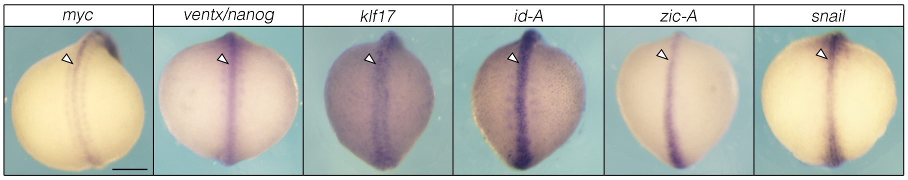


**Extended Data Fig. S4.** Neural crest and pluripotency factors are expressed along the antero-posterior axis in lamprey embryos. Scale bar: 250 µm.

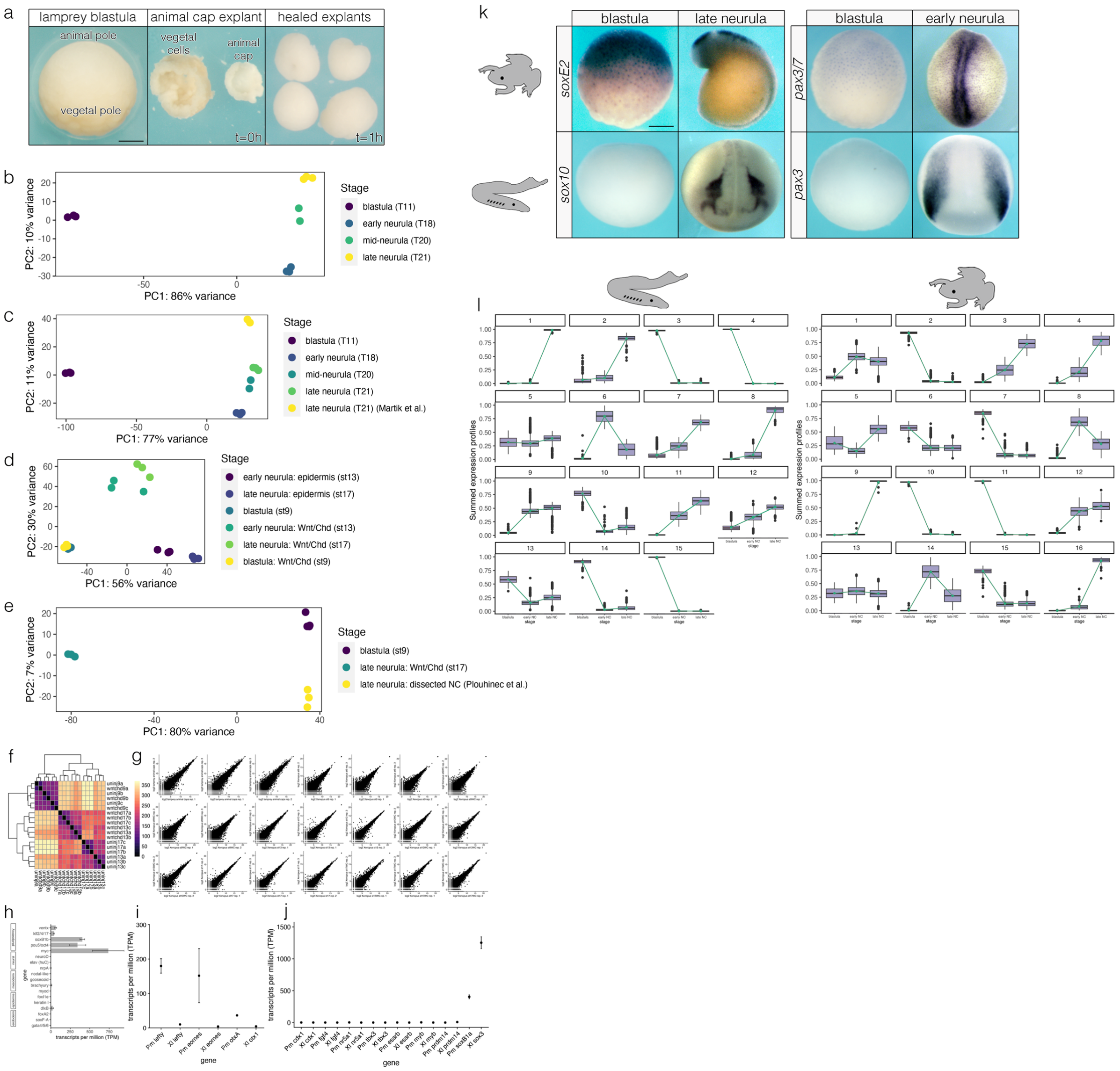


**Extended Data Fig. S5**. Analysis of RNA-Seq data in *Xenopus* and lamprey. (a) Example dissection of lamprey animal caps. (b-e) Comparisons of principal components analyses (PCA) of samples used in this study and elsewhere indicate a lack of significant batch effects. (f,g) Reproducibility of RNA-Seq data generated in this study for *Xenopus*. (h) Lamprey animal caps express canonical pluripotency factors but not markers of germ layer differentiation. (i) Genes involved in vertebrate pluripotency that are expressed at in blastula-stage lamprey embryos are not expressed in *Xenopus* embryos blastulae. (j) Several factors involved in mammalian pluripotency are absent in the animal poles of *Xenopus* and lamprey. Transcript abundance of *soxB1* factors are included for both *Xenopus* (*sox3*) and lamprey (*soxB1a*) for context. Pm = *Petromyzon marinus*, Xl = *Xenopus laevis*. (k) Validation of RNA-Seq comparisons showing species-specific differences in expression of neural crest genes in the blastula. Lamprey *soxE2* and *pax3/7* are expressed in both animal pole cells and neural crest, whereas these factors are only enriched in the neural crest of *Xenopus.* (l) Full results of *k*-means analysis. Abbreviations: uninj9 = uninjected caps, st9; wntchd9 = wnt8a/chordin-injected caps, st9; uninj13 = uninjected caps, st13; wntchd13 = wnt8a/chordin-injected caps, st13; uninj17 = uninjected caps, st17; wntchd17 = wnt8a/chordin-injected caps, st17. Scale bars: 250 µm.

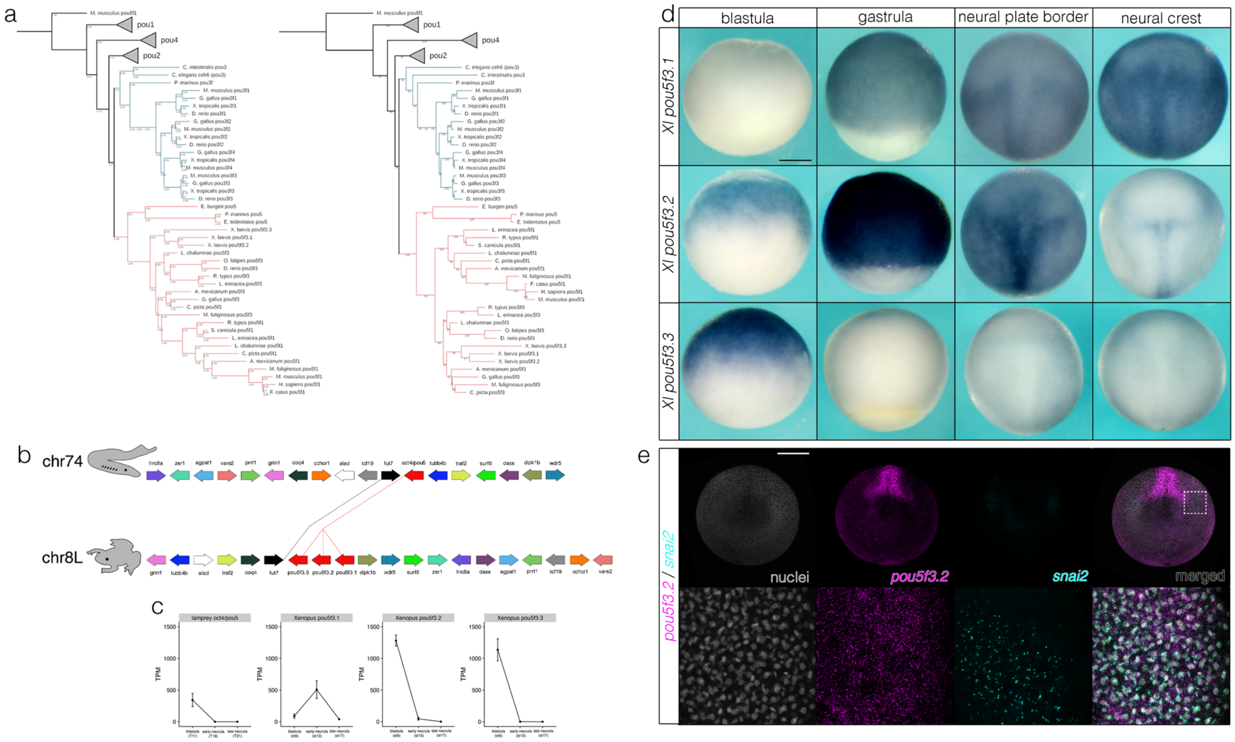


**Extended Data Fig S6.** Phylogenetic analysis and characterization of lamprey *pou5* and *Xenopus pou5f3*. (a) Bayesian (left) and Maximum likelihood (right) phylogenies of POU transcription factors. (b) Synteny comparisons of *pou5* loci. (c) Quantification of transcript abundance (TPM) by RSEM for lamprey *pou5* and *Xenopus pou5f3* in animal pole cells and neural crest. Data were obtained from ≥100 lamprey animal caps for n = 3 biological replicates and ≥10 Xenopus animal caps at each stage for n = 3 biological replicates. (d) *in situ* hybridization of *Xenopus pou5f3* paralogs. (e) HCR-FISH showing colocalization of *pou5f3.2* with *snai2* transcripts in the neural plate border. Xl, *Xenopus laevis*. Pm, *Petromyzon marinus*. Data were obtained from n = 3 biological replicates. Scale bars: 250 µm.

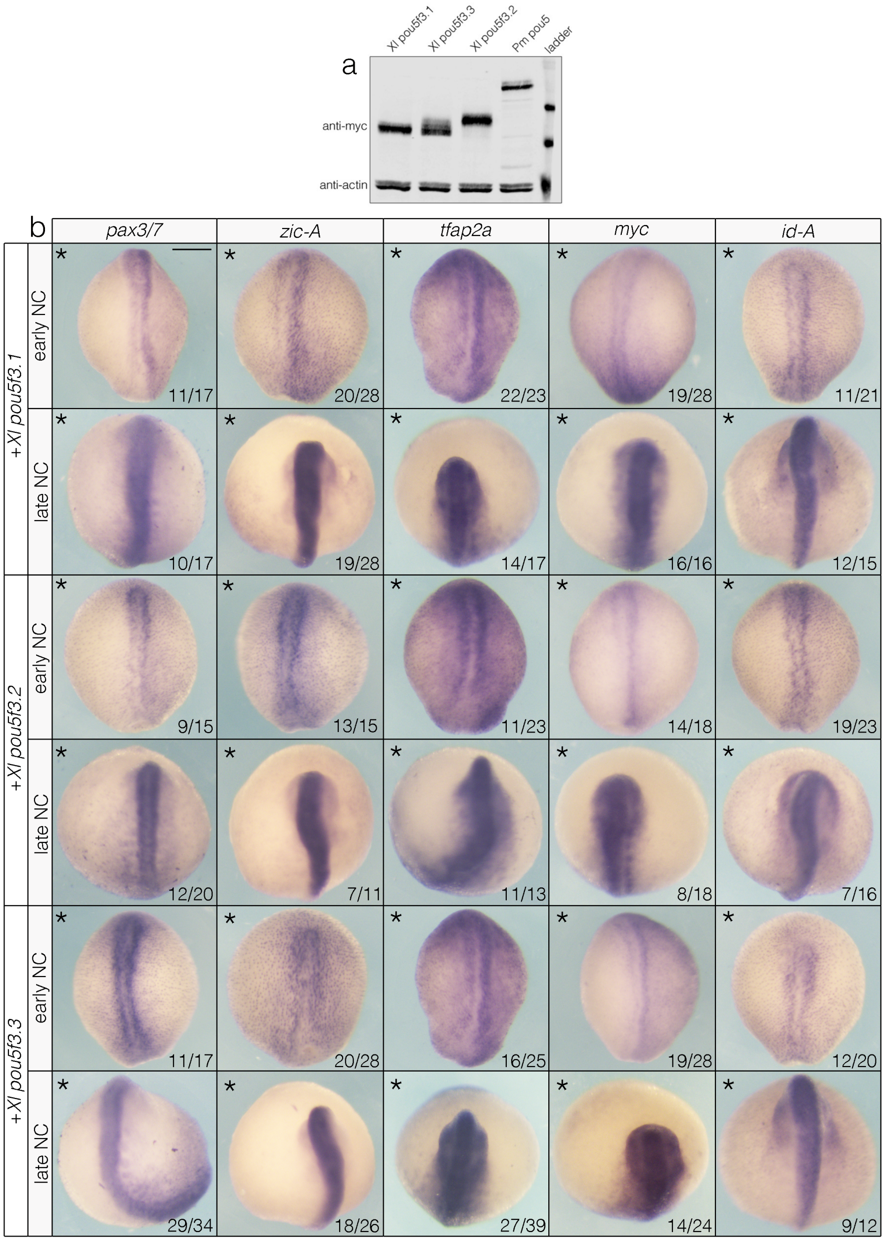


**Extended Data Fig. S7.** *Pou5f3* overexpression experiments in lamprey embryos. (a) Western blot showing matched protein levels for experiments. (b) Overexpression of *Xenopus pou5f3* factors in lamprey causes loss or no change of expression of multiple neural plate border and neural crest markers. Asterisk denotes the injected side of the embryo. Scale bar: 250 μm.

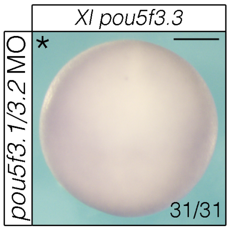


**Extended Data Fig. S8**. MO knockdown of *Xenopus pou5f3.1* and *pou5f3.2* does not result in a compensatory increase in *pou5f3.3* expression. Scale bar: 250 µm.

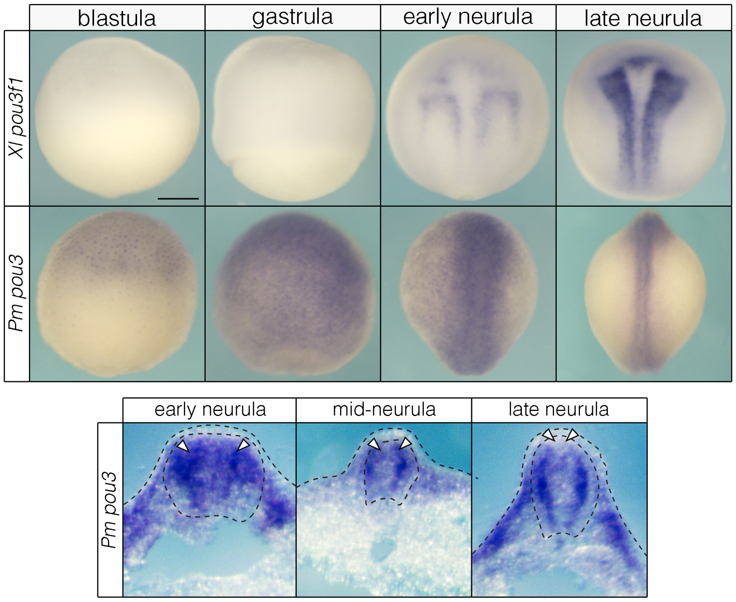


**Extended Data Fig. S9.** Whole mount (top) and sectioned (bottom) embryos showing wildtype *pou3* expression in *Xenopus* (Xl) and lamprey (Pm) embryos. In lamprey, *pou3* transcripts are observed in the neural crest of early and mid-neurulae (arrowheads). By late neurula stages, *pou3* expression is largely absent from the neural crest (arrowheads) while being enriched in the neural tube. Scale bar: 250 µm.

### Descriptions of Extended Data Source Files

**Extended Data Source File S1**: spreadsheet associated with Fig. 3b containing log transformed TPMs used for correlation analyses.

**Extended Data Source File S2**: spreadsheet associated with Fig. 3d containing output from DESeq2 analysis for *Xenopus* and lamprey.
